# Age associations with cortical and subcortical brain structure in adolescents age 9-17

**DOI:** 10.1101/2025.09.18.677236

**Authors:** Diana M. Smith, Alison Rigby, Diliana Pecheva, Pravesh Parekh, Ashley Becker, Robert Loughnan, Thomas E. Nichols, Terry L. Jernigan, Anders M. Dale

## Abstract

**Introduction:** Adolescence is a pivotal period in brain structural development and maturation. However, investigation of cortical and subcortical brain changes during this time have been limited by small sample size and have generally examined the brain at the level of predetermined regions of interest. The recently developed Fast Efficient Mixed-Effects Algorithm (FEMA) allows for increased computational speed using mixed-effects models applied at the voxel or vertex level, as well as across multiple regions of interest.

**Methods:** We extended the existing FEMA framework to represent predictors using natural spline basis functions, enabling us to model nonlinear trajectories of brain structure as a function of age. We then applied this model to the The Adolescent Brain Cognitive Development℠ Study (22,651 observations from 10,521 unique subjects aged 9.00-17.77) to study the age-related trajectories of tabulated cortical and subcortical volumes, vertexwise cortical thickness and surface area, and voxelwise volume assessed using the Jacobian. Models are reported separately in males and females.

**Results:** Global volume variables, including total subcortical gray matter volume, peaked near 13 years in females and 15 years in males. Vertexwise cortical surface area followed an inverted U-shaped curve, whereas vertexwise cortical thickness followed a monotonic decrease during the age range studied. Voxelwise imaging analysis revealed regional differences in age trajectories at the subregional level.

**Discussion:** The results of this work replicate and extend prior findings related to adolescent brain development, and illustrate distinct spatiotemporal patterns of structural changes in subcortical regions. The updated FEMA framework is publicly available for use in similar large datasets.

## Introduction

Brain development during childhood and adolescence is associated with distributed structural alterations in both gray matter (GM) and white matter (WM) that occur concurrently with cognitive and behavioral development (Palmer and Jernigan 2023). Cortical thickness decreases in a nearly linear fashion during adolescence, whereas cortical surface area peaks in early adolescence and then decreases during the teen years, with region-specific differences in timing across the cortical sheet (Jernigan, Trauner, et al. 1991; Jernigan et al. 2016).

Subcortical GM volumes have been studied less but are also thought to decrease with age during adolescence (Jernigan, Trauner, et al. 1991; Pomponio et al. 2020). However, prior studies have been mainly in small samples with dozens or hundreds of participants, and the use of linear and quadratic models for this application has been questioned (Walhovd et al. 2016), leading to the use of spline functions for the modeling of curvilinear brain imaging trajectories (Bethlehem et al. 2022). This study presents the systematic investigation of age-related trajectories of brain morphometry, using a combination of regions-of-interest (ROIs), vertexwise, and voxelwise data to better understand the distributed spatiotemporal patterns in brain volumes.

Global measures of cortical surface area (CSA) and apparent cortical thickness (CTH) have unique development trajectories over adolescence and have been correlated with cognitive measures (Palmer 2021). CSA expands during early childhood, then slowly contracts over adolescence (Jernigan et al. 2016; Bethlehem et al. 2022). Conversely, prior research has found that CTH decreases at a consistent rate from early childhood through adolescence (Brown et al. 2012; Jernigan et al. 2016; Bethlehem et al. 2022). However, from the very first observations of decreasing cortical thickness with age by Jernigan and colleagues (1991; 1991), there has remained the possibility that “apparent” cortical thinning observed on MRI could also be due, in part, to increased myelination.

GM volume trajectory differences in the frontal, parietal, and temporal lobes have been observed over adolescence, with a more pronounced difference between GM and WM volume in the frontal lobe over this period (Matsuzawa 2001). Fronto-temporal GM volume peaks later and declines at a slower rate than primary sensory regions (Giedd 2004; Bethlehem et al. 2022). Region-specific trajectories in CSA and CTH during adolescence, which are both associated with cortical GM volume, may have specific relationships to cognition, as measured by performance on cognitive tasks. For example, anterior cingulate gyrus CSA has been associated with reaction time on the NIH Cognitive Toolbox Flanker task in adolescents (Fjell et al. 2012; Jernigan et al. 2016) and inferior fronto-parietal CSA was associated with the Stop Signal Reaction Time in children (Curley et al. 2018).

Vertexwise analyses have revealed the relative gaps left by regional investigations. Palmer and colleagues (2021) found that so-called “fluid” and “crystallized” domains of cognitive function were associated with distinct regional patterns in CSA and CTH. Of note, *not* controlling for sociodemographic factors increased the similarity between these regionalization patterns, highlighting the importance of sociodemographic factors in the relationship between brain structure and performance on standardized cognitive tasks. Additionally, this study emphasized the value of high-dimensional vertexwise data in illuminating patterns that are not easily visualized through regional analyses alone.

In a large meta-analytic analysis from Bethlehem and colleagues (2022), total GM volume was observed to peak at age 6, then decreases for the rest of childhood and adolescence. The specific timing of subcortical structural change has been found to vary across nuclei, with one study finding the strongest negative effect of age on thalamus, caudate, and lenticular nucleus volume with smaller relationships to the nucleus accumbens and basomesial diencephalon (Sowell et al. 2002). One recent study found strong age associations with basal ganglia microstructure, estimated using restriction spectrum imaging (Palmer et al. 2022), though to our knowledge, there has been no investigation into the volume trajectories of these and other subcortical regions in a large longitudinal adolescent sample.

Sex differences are likely to influence age-related trajectories of some subcortical volumes (Sowell et al. 2002). Giedd and colleagues (1997) reported that amygdala volume increases more in males, whereas the hippocampus increases more in females, and suggested that these difference may be related to the high densities of sex steroid hormone receptors in the medial temporal lobe and the trophic effect of sex steroids on these structures. For this reason, it is common practice to estimate age-related trajectories in males and females separately in recent research (Bodison et al. 2020).

### Limitations of prior research

The 2000s witnessed several studies of age associations with cortical thickness, which yielded mixed results (reviewed in Walhovd et al. 2016). One possible explanation for this heterogeneity is related to low statistical power in many studies, which allows for sample heterogeneity and cohort effects. Button and colleagues (2013) discussed the impact of underpowered, low dimensional studies on replicability in neuroscience research, where it is likely that many outcomes are driven by multiple small effects. Jernigan and colleagues (2016) called for large-scale longitudinal datasets with comprehensive measurement of potential environmental and experiential mediators that may influence developmental trajectories. The Adolescent Brain Cognitive Development^℠^ Study (ABCD Study^®^) was designed to provide one such dataset (Volkow et al. 2018).

Another critique of early investigations into age trajectories of structural morphometry focused on the use of quadratic models to account for age effects, which can lead to differing estimates of when the “peak” occurs based on the range of ages assessed (Walhovd et al. 2016). This has led to a recommendation in the field to use other methods, such as splines (piecewise polynomial functions), rather than quadratic models, as splines have been observed to outperform quadratic models when fit to brain imaging data across different age ranges (Fjell et al. 2010). Indeed, in the past decade, there has been a shift toward using smooth functions such as splines (piecewise polynomials) when modeling age (e.g., Bethlehem et al. 2022).

Nonlinear models have been primarily employed to document the trajectories of global brain imaging variables such as GM and WM volume, as well as region-wise measurements of apparent cortical thickness and surface area. For example, in a sample of 2,000 infants and children age 0-6 years, Alex and colleagues (2024) observed that the amygdala was the first subcortical volume to reach 99% of its asymptotic volume according to a nonlinear mixed model using an asymptotic function; other subcortical structures achieved this milestone at varying ages, under the same model parameters. One analysis harmonized structural brain MRI data from over 10,000 participants across eighteen studies to model age trajectories of individual cortical and subcortical ROIs from ages 3 to 96 (Pomponio et al. 2020). Results showed relative peaks in subcortical volumes during adolescence, with heterogeneity across ROIs regarding the precise timing of the peak volume. However, the harmonization of several distinct samples remained one limitation of this study, which included only two datasets (1,750 participants) with data collected prior to age 16. The largest study to date was performed by Bethlehem and colleagues (2022), who found nonlinear decreases in global GM volume during adolescence, with frontotemporal regions peaking in volume later and with a slower decline in volume compared to sensorimotor cortices. Few studies have examined age associations with cortical morphometry in a vertexwise fashion, and to our knowledge, none have investigated age variation in voxelwise structural metrics.

### The current study

Accordingly, we applied a generalized additive mixed-effects model using natural cubic splines to characterize the age-related trajectories of cortical and subcortical structural brain morphometry in a whole-brain manner. We employed this model across regional measurements of subcortical volumes, as well as vertexwise cortical thickness and surface area. Finally, we applied the model to the Jacobian determinant to achieve granular estimates of the associations between age and relative volume changes over time.

## Methods

### Sample

The ABCD Study^®^ consists of 11,880 adolescents across 21 sites, beginning at age 9-11 years and with longitudinal study visits occurring regularly (with imaging visits occurring every two years; Volkow et al. 2018). This paper uses data from the ABCD Study^®^ 6.0 Data Release. The ABCD Study^®^ cohort is epidemiologically informed and the sample has been curated to closely reflect the adolescent population of the United States (Garavan et al. 2018). Eligibility criteria for the ABCD Study^®^ included a) English proficiency in the child, b) the absence of severe sensory, neurological, medical or intellectual limitations that would inhibit the child’s ability to complete study protocol, and c) inability to complete a MRI scan at the baseline visit. All study protocols were approved by the University of California, San Diego Institutional Review Board. Parent or caregiver permission and youth assent were obtained for each participant.

Table 1 provides demographic information for the analytic sample. Statistical analyses included 22,651 observations from 10,521 unique subjects. 7,073 subjects had data from at least two timepoints. Participants aged 9.00 - 17.77 years were included in analysis. Observations were included in the final sample if the participant had complete data across sociodemographic factors (household income and parental education), genetic data, available imaging data that passed all inclusion criteria, and available information regarding acquisition scanner ID and software version.

**Table 1.** Demographics of the sample.

|  | Baseline | 2 Year FU | 4 Year FU | 6 Year FU | *p* |
| --- | --- | --- | --- | --- | --- |
| **n** | 9009 | 6693 | 5221 | 2183 |  |
| **interview_age (mean (SD))** | 9.97 (0.63) | 12.00 (0.65) | 14.17 (0.73) | 16.11 (0.65) | <0.001 |
| **sex (%)** |  |  |  |  | 0.191 |
| **F** | 4388 (48.7) | 3114 (46.5) | 2458 (47.1) | 1045 (47.9) |  |
| **I** | 2 (0.0) | 1 (0.0) | 1 (0.0) | 0 (0.0) |  |
| **M** | 4619 (51.3) | 3578 (53.5) | 2762 (52.9) | 1138 (52.1) |  |
| **high.educ (%)** |  |  |  |  | <0.001 |
| < HS Diploma | 448 (5.0) | 311 (4.6) | 260 (5.0) | 87 (4.0) |  |
| Bachelor | 2687 (29.8) | 2022 (30.2) | 1521 (29.1) | 700 (32.1) |  |
| HS Diploma/GED | 830 (9.2) | 604 (9.0) | 458 (8.8) | 170 (7.8) |  |
| Post Graduate Degree | 2424 (26.9) | 1843 (27.5) | 1554 (29.8) | 684 (31.3) |  |
| Some College | 2620 (29.1) | 1913 (28.6) | 1428 (27.4) | 542 (24.8) |  |
| **pds_y_p_average (mean (SD))** | 1.89 (0.76) | 2.60 (0.95) | 3.61 (0.76) | 4.11 (0.58) | <0.001 |
| **anthro_bmi_corr (mean (SD))** | 18.66 (4.01) | 20.60 (4.79) | 22.75 (5.42) | 23.56 (5.25) | <0.001 |
| **demo_comb_income_v2_l (%)** |  |  |  |  | <0.001 |
| Less than 5,000 | 301 (3.3) | 179 (2.7) | 110 (2.1) | 32 (1.5) |  |
| \$5,000 through \$11,999 | 302 (3.4) | 187 (2.8) | 128 (2.5) | 35 (1.6) | |
| \$12,000 through \$15,999 | 222 (2.5) | 126 (1.9) | 92 (1.8) | 30 (1.4) | |
| \$16,000 through \$24,999 | 409 (4.5) | 233 (3.5) | 185 (3.5) | 47 (2.2) | |
| \$25,000 through \$34,999 | 537 (6.0) | 367 (5.5) | 275 (5.3) | 88 (4.0) | |
| \$35,000 through \$49,999 | 760 (8.4) | 488 (7.3) | 355 (6.8) | 119 (5.5) | |
| \$50,000 through \$74,999 | 1231 (13.7) | 927 (13.9) | 599 (11.5) | 186 (8.5) | |
| \$75,000 through \$99,999 | 1317 (14.6) | 957 (14.3) | 697 (13.3) | 288 (13.2) | |
| \$100,000 through \$199,999 | 2859 (31.7) | 2308 (34.5) | 1837 (35.2) | 857 (39.3) | |
| \$200,000 and greater | 1071 (11.9) | 921 (13.8) | 943 (18.1) | 501 (23.0) | |

Note. Demographic data is shown for age in years (mean (SD)), sex at birth, household income, parental education, mean of youth- and parent-reported pubertal status, and body mass index. Data are stratified by timepoint: baseline, 2-year, 4-year, and 6-year follow-up. There were significant differences in age, income, education, pubertal status (average of parent and youth report), and BMI across timepoints. Variable names from the tabulated data release, as well as variable names corresponding to the provided code, are used in the table for replication.

### MRI acquisition

Imaging data for the ABCD Study^®^ were collected at all 21 sites, using Siemens Prisma, GE 750 and Philips Achieva and Ingenia 3T scanners. The structural and diffusion imaging acquisition protocols and image processing pipeline used in the ABCD Study^®^ have been described elsewhere (Casey et al. 2018; Hagler et al. 2019). Briefly, T1-weighted images were acquired using a 3D magnetization-prepared rapid acquisition gradient echo (MPRAGE) scan with 1mm isotropic resolution and no multiband acceleration. T1w structural images were corrected for gradient nonlinearity distortions using scanner-specific, nonlinear transformations provided by scanner manufacturers (Wald et al. 2001; Jovicich et al. 2006). Intensity inhomogeneity correction was performed by applying smoothly varying, estimated B1-bias fields to the T1w intensities, using a standard target white matter intensity value (Hagler et al. 2019). Modalities were aligned within each participant using the rigid body transforms from the ABCD processing pipeline (Hagler et al. 2019).

### Atlas registration and Jacobian

Images were registered to the ABCD Study^®^ atlas generated using the Multimodal Image Normalisation Tool (MINT; Pecheva et al. 2022). Registration was based on eleven input channels: 3D T1, zeroth and second order spherical harmonic (SH) coefficients from the restricted fiber orientation distribution (FOD), zeroth order SH coefficient from the hindered and free water FODs, WM, and GM segmentations. The registration target was the group mean image, constructed iteratively, and transformation fields were estimated for all participants towards the provisional group mean. Participants with poor registration to the atlas, defined as a mean voxelwise correlation to atlas of <0.8, were excluded from statistical analysis.

The differences in tissue volume between each participant’s scan and the atlas can be described using the nonlinear warps from the image registration process. We calculated the partial spatial derivatives of the nonlinear deformation field at each voxel, the Jacobian matrix, and its determinant. After global scaling, the Jacobian determinant at the voxel-level represents the local scaling factor of volume under a transformation, indicating expansion (determinant > 1) or contraction (determinant <1). Of note, the Jacobian determinant is calculated after global scaling in the x, y and z directions, based on affine registration to atlas space, therefore this metric represents relative volume variability rather than absolute volume variability.

### Labeling regions of interest

ROI-wise analysis of subcortical volumes used the volume estimates from the Freesurfer 5.3 segmentation (Fischl et al. 2002). These data are readily available in tabulated form in the ABCD Study^®^ 6.0 data release and represent a summary of the native space voxelwise data for each participant. A full list of tabulated volume variables used in the primary analysis is available at the online ABCD Study^®^ data dictionary: https://nbdc-datashare.lassoinformatics.com/data-wrangler, table name “mr_y_smri vol aseg”. Additional subcortical nuclei, not available in the Freesurfer segmentation, were labeled by registering readily available high spatial resolution atlases to our atlas space. Bilateral binary masks were created for each ROI by thresholding at 0.8 probability across the ROI, meaning that in a given voxel, at least 80% of participants showed that ROI label. The Najdenovska thalamic nuclei atlas was generated using a k-means algorithm, taking as inputs the mean FOD SH coefficients from within a Freesurfer parcellation of the thalamus using data from 70 HCP subjects (Najdenovska et al. 2018). Interpolated binary ROI masks were created for all ROIs from both the Pauli and Najdenovska atlases.

### Statistical analysis

Univariate general additive mixed effects models (GAMMs) were applied to each ROI (using the tabulated volume data as the dependent variable), vertex (for cortical thickness and cortical surface area), or voxel (for the voxelwise Jacobian data). All of the main results shown are from the model depicted below with a smooth function of age included as a single predictor in long format and the longitudinal component modelled as a random effect of subject. This model was applied to the full sample using a fixed effect of sex, as well as in sex-stratified analyses; because a main effect of sex was observed in nearly all models, sex-stratified analyses are presented in this work. Given the demographic diversity in the sample, all statistical analyses controlled for the sociodemographic variables household income and parental education.The top 10 genetic principal components were used to account for ancestry effects in lieu of self-declared race/ethnicity. Additional fixed effects included scanner ID and MRI software version.

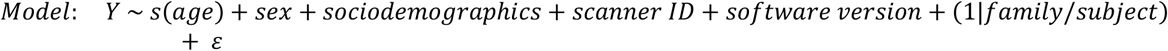

To construct the GAMM, we first calculated a basis matrix for natural cubic splines with unit heights at knots using the *createBasisFunctions* function from the *cmig_tools* package in MATLAB (Parekh et al. 2023). The bases were applied to an age range of 9 to 18, with four evenly spaced knots: two boundary knots and two internal knots. The resulting basis matrix was demeaned to avoid redundancy with the intercept term in the model. This demeaning step resulted in a rank deficient matrix, so the middle basis function was dropped to achieve full rank. After demeaning to avoid redundancy with the intercept term, we applied singular value decomposition to the resulting basis matrix, which orthogonalizes the bases and ensures independence by removing redundant bases. The remaining bases were rescaled to values between 0 and 1 and interpolated with the age data, results of which were then passed to a linear mixed-effects model using Fast Efficient Mixed-Effects Algorithm, a fast mixed-effects modeling tool that was developed for the ABCD Study and similar datasets (Parekh et al. 2023). FEMA computed regression coefficients for each basis function, which were then linearly combined to construct the best-fit spline function for age. Compared to smoothing splines, regression splines incorporate a smaller number of internal knots and do not introduce a penalty (Perperoglou et al. 2019); we therefore did not penalize the resultant spline function.

To aid in interpretation of estimated spline functions, all covariates were mean-centered. Categorical variables (MRI device number and software version, household income) were releveled so that the most common value was used as the reference value. As a result, graphs of the model-estimated outcome as a function of age represent the estimated morphometric measurement for a hypothetical participant with average values for all continuous covariates, with the most common value for categorical variates, averaging across sex.

We estimated 95% confidence intervals by first estimating the variance of the function as:

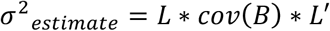

Where *L* represents a zero-padded matrix of basis function values plus the value of the intercept for each age, *cov*(*B*)represents the covariance matrix of the regression coefficients as estimated by FEMA (Liu 2015). We then multiplied the square root of this variance estimate by 1.96 to get the upper and lower ranges of the confidence interval.

Unthresholded maps of the beta estimate for s(age) are presented in the main figures to provide an estimate of the annualized rate of change, calculated as the derivative of the smooth function at a given age. The unthresholded maps were chosen to provide a comprehensive description of the continuous distribution of effects, rather than displaying only effects that pass a conservative Bonferroni correction. Animations of age associations with cortical thickness, cortical surface area, and voxelwise JA were prepared using FIJI using a frame rate of 8 fps (125 ms per frame; Schindelin et al. 2012).

Data preprocessing was completed in R version 4.2.3 as well as MATLAB v2022a. Statistical analyses were conducted using MATLAB v2022a. The code for FEMA is publicly available at https://github.com/cmig-research-group/cmig_tools and the code used for this project is available at https://github.com/dmysmith/age.

## Results

### Age effects on tabulated cortical and subcortical volumes

We calculated the model-estimated subcortical volume for each region of interest using the Desikan-Killiany parcellation for subcortical structures. Supplementary Figure 1 shows the estimated volumes for each structure as a function of age. With the exception of WM hypointensities at the earlier portion of the age range evaluated (smri_vol_scs_wmhint), there was a main effect of sex where males had larger volumes of all ROIs investigated than females.

In trajectories of subcortical gray volume (smri_vol_scs_subcorticalgv), females exhibited increases in volume estimates at the beginning of the age range, followed by a peak between 12 and 13 years, then sharper decreases afterwards; in contrast, males peaked later, at approximately 15 years (Supplementary Figure 1). Similarly, for whole brain volume (smri_vol_scs_wholeb) and supratentorial volume (smri_vol_scs_suprateialv), females peaked between age 11 and 12, whereas males peaked around age 13 and displayed more gradual decreases thereafter. The total cerebellar cortical volume (smri_vol_scs_crbcortexlh, smri_vol_scs_crbcortexrh) also peaked in females near age 12, whereas males peaked around age 14. Total cerebellar WM volume (smri_vol_scs_crbwmatterlh, smri_vol_scs_crbwmatterrh) followed a similar pattern but with more protracted trajectories compared to the cerebellar cortical volume (cerebellar WM peaked at 13 years in females and plateaued at 15 years in males).

Of the subcortical ROIs, the anterior corpus callosum (smri_vol_scs_ccat) was noted to exhibit markedly different trajectories between females and males, where females followed a steeper trajectory prior to age 13 followed by slower increases at subsequent ages, whereas males exhibited an increased slope of the age curve from age 13 to 16.

Within the subcortical ROIs, the amygdala (smri_vol_scs_amygdalalh, smri_vol_scs_amygdalarh) and putamen (smri_vol_scs_putamenlh, smri_vol_scs_putamenrh) both displayed peak volumes near age 12 in females (Supplementary Figure 1). Males exhibited a later peak in amygdala volume near age 14 (in the left hemisphere) and 16 (in the right hemisphere), followed by gradual decreases in volume. In the left thalamus (smri_vol_scs_tplh), males exhibited steeper increases in volume until age 14, when increases became more gradual; on the other hand, in the right thalamus (smri_vol_scs_tprh), the model estimated a slight decrease in volume starting at age 15 in males. Compared to the amygdala and putamen, the hippocampus and pallidum appeared to have more protracted trajectories over the course of the assessed age range, with peak volumes occurring in females around 13 years and males around 14-16 years (Supplementary Figure 1). Supplementary Figure 2 and 3 depict cortical surface area and cortical thickness according to APARC ROIs, with each dependent variable estimated as a smooth function of age.

### Age effects on vertexwise cortical thickness and cortical surface area

Figure 1 depicts the vertexwise slope of the model-estimated cortical surface area as a function of age. In general, there was a global pattern of increasing surface area with age until approximately age 11-12, followed by global decreases in vertexwise surface area after that point. There were regional differences in exact timing of this peak in surface area, with more parts of the occipital and parietal cortices beginning to decrease earlier and the most protracted trajectories occurring in the medial frontal and temporal cortices (Figure 1). By age 15, most of the decreases in surface area were slower, except for the orbitofrontal cortex, which continued to decrease in surface area through the end of the age range. Standard error of model-estimated surface area is depicted in Supplementary Figure 4.

**Figure 1.**
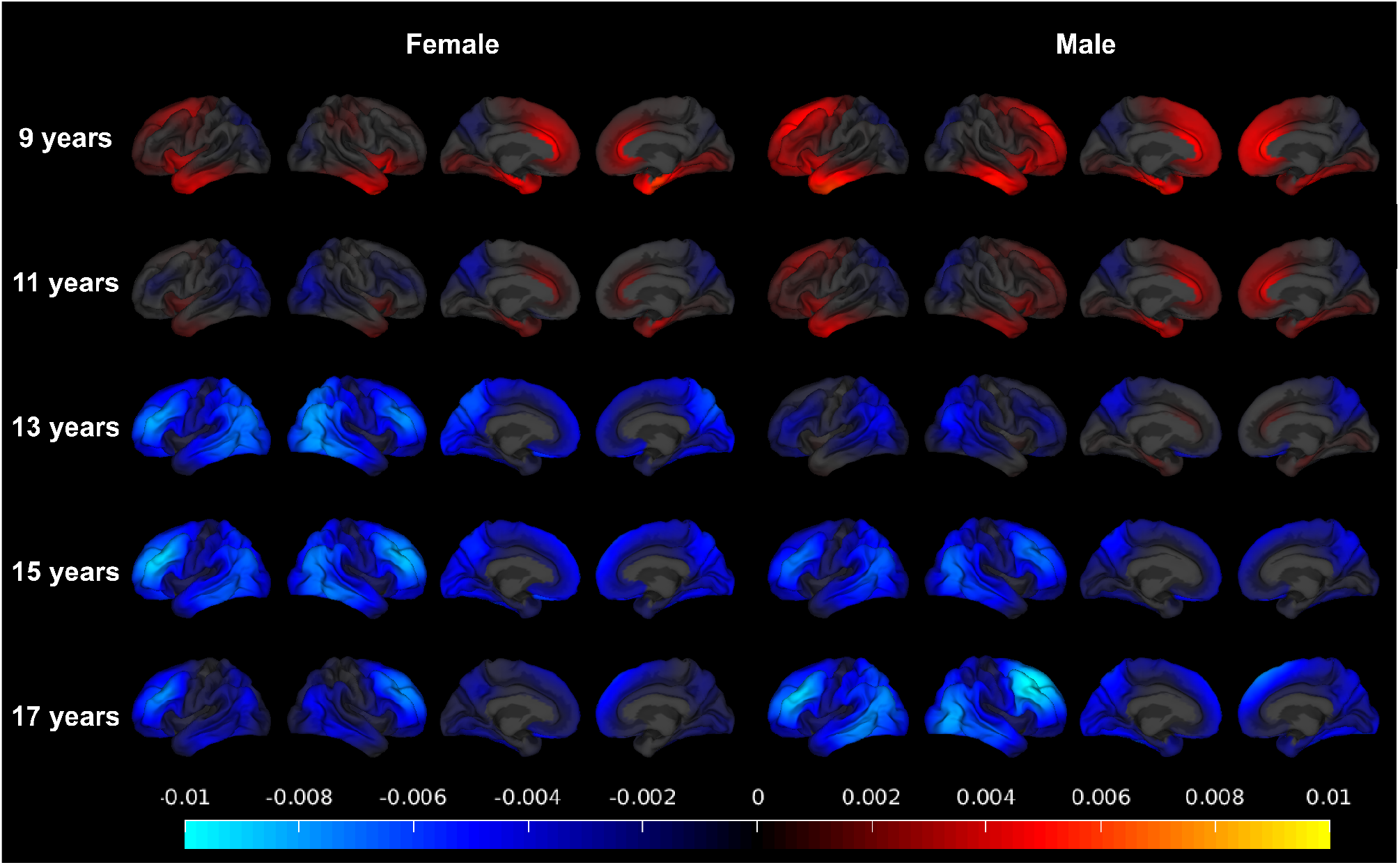
Vertexwise plots of the derivative of model-estimated cortical surface area as a function of age, estimated separately in females (left four columns) and males (right four columns). Estimates are represented as beta estimates in units of square millimeters per month of age.

Vertexwise cortical thickness displayed decreases throughout adolescence (Figure 2). The steepest decreases occurring in early adolescence (ages 9-11) were observed in the medial frontal cortex, with the medial parietal cortex exhibiting similar decreases in thickness from ages 11-13. Later in adolescence (age 13) the most pronounced changes were observed in the medial frontal and orbitofrontal cortices (Figure 2).

**Figure 2.**
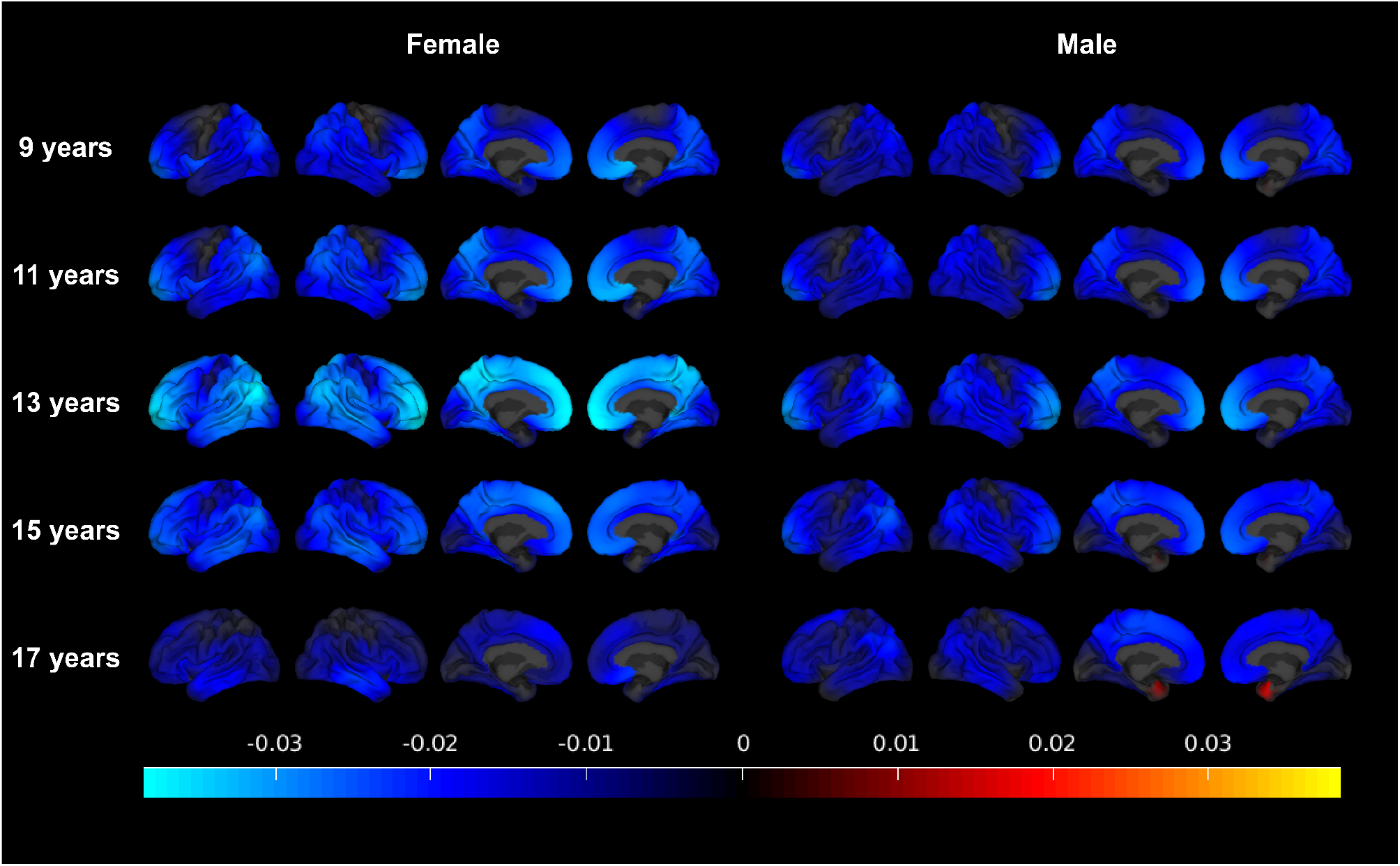
Vertexwise plots of the derivative of model-estimated apparent cortical thickness as a function of age, estimated separately in females (left four columns) and males (right four columns). Estimates are represented as beta estimates in units of millimeters per month of age.

### Age effects on voxelwise Jacobian

Figure 3 depicts the voxelwise slope of the model-estimated Jacobian as a function of age. Age effects followed a distributed pattern consistent with anatomical structures, with positive age effects in the pallidum and negative age effects in the caudate, putamen, and parts of the thalamus. WM tracts showed a positive relationship with age that increased toward the latter portion of the evaluated age range, whereas cortical GM exhibited a negative relationship with age that also increased in magnitude over the course of the age range (Figure 3).

**Figure 3.**
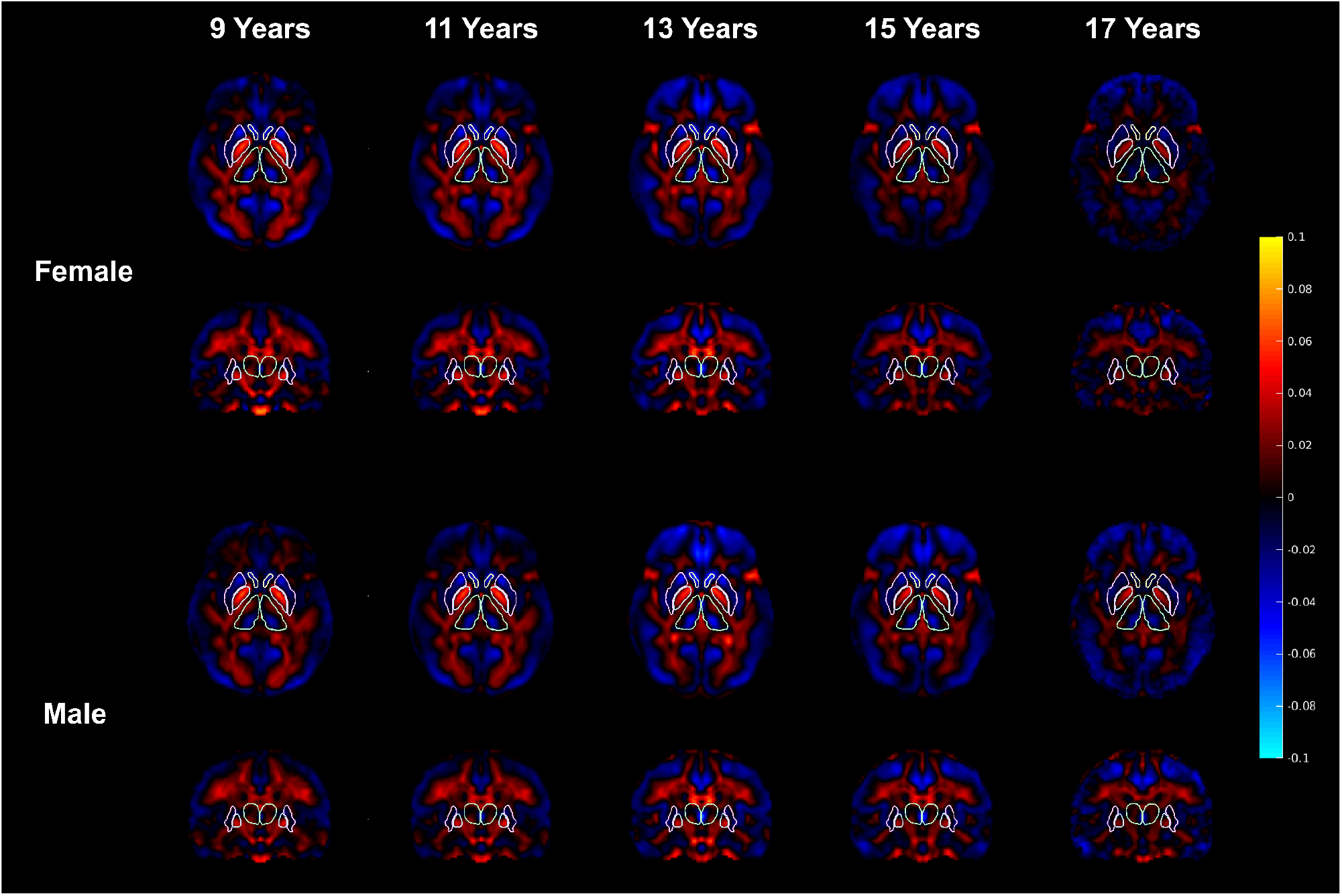
Voxelwise plots of the derivative of model-estimated Jacobian as a function of age, estimated separately in females (top two rows) and males (bottom two rows). Beta estimates are depicted, representing the change in standardized units of the Jacobian per year of age. For ease of interpretation, outlines of the accumbens area, putamen, pallidum, and thalamus are provided from the ABCD3 MINT atlas (Pecheva et al. 2022).

## Discussion

This study investigated age associations with tabulated, vertexwise, and voxelwise morphometric measurements in the ABCD Study^®^. The trajectories observed within the tabulated subcortical volumes followed similar patterns to those observed in prior research, allowing for more powerful estimation using a large longitudinal sample. Vertexwise cortical surface area followed an inverted U-shaped curve similar to that noted in prior work (Jernigan et al. 2016), and vertexwise cortical thickness followed a generally monotonic decrease (Palmer and Jernigan 2023). Voxelwise estimation of the Jacobian as a function of age provided a higher resolution of effects, which illuminated anatomical distributions of age trajectories at a more granular level compared to regional analyses.

Global volume variables, including total subcortical GM volume, were found to peak near 13 years in females and 15 years in males. This finding extends those from recent meta-analyses of brain imaging studies (Pomponio et al. 2020; Bethlehem et al. 2022), and enhances fine details regarding precise timing of adolescent trajectories. For example, the LIFESPAN dataset (Pomponio et al. 2020) estimates an increase in total brain volumes in males throughout the teenage years, whereas our larger sample with several timepoints throughout adolescence estimated a peak in whole brain volume near age 13.

The tabulated volume results shed light on Inconsistencies in the literature regarding sexual dimorphism in the increase in WM volume seen during adolescence (Peper et al. 2011). We found that global variables related to WM (e.g., total cerebral WM, total cerebellar WM, and corpus callosum) exhibited different trajectories between males and females: females tended to display steeper increases earlier in adolescence, whereas males displayed more protracted increases throughout the age range studied. With this work, we leverage the large ABCD Study^®^ sample to chart specific trajectories with narrow confidence intervals, allowing for the more detailed illustration of these sex-specific processes.

Subcortical GM volume metrics displayed overall decreases during adolescence, extending findings from smaller studies of subcortical volume. Our findings are similar to previous studies with smaller sample sizes, which found heterogeneity across GM and WM structures, including subcortical volumes across adolescence and early adulthood (Sowell et al. 2002; Østby et al. 2009). Vertexwise models of age associations with cortical thickness and cortical surface area provide a more precise view of the timing of these developmental changes across the cortical sheet. In general, our findings provide insight into the wave of structural changes occurring at different timepoints across the cortex, which deepens our understanding of the temporal dynamics of cortical maturation. One of the major aims of the ABCD Study^®^ was to establish more meaningful growth curves for adolescent brain development (Volkow et al. 2018); this work provides a direct contribution to that aim.

The incorporation of the voxelwise Jacobian is a novel method for measuring structural brain morphometry at the voxel level. Of note, the Jacobian takes into account the total x, y, and z scaling for each participant scan, making this measurement more similar to a total volume-adjusted volume metric, more suited to addressing questions of *relative* regional growth rather than absolute volume. In other words, our findings using the Jacobian found that cortical GM was largely decreasing after age 12 (with the exception of the frontal cortices), and WM tracts began to display positive age associations after age 12.

The current study should be interpreted in light of several limitations. First, there is ongoing discussion about the comparative utility of GAMMs (such as the model used in this work) compared to other advanced regression models. Multivariable Fractional Polynomial Regression has been found in one recent comparative analysis to outperform a generalized additive model implemented in a large sample of brain imaging data (Ge 2023). Second, it should be noted that the Jacobian metric used in this work is calculated after having been log-transformed, which may hinder the intuitive interpretation of effect sizes. Third, while the spline function implemented in our work has been demonstrated to result in better model fits than a simple quadratic model (Walhovd et al. 2016), it is often the case that using a longitudinal sample spanning many timepoints over a large age range is the most advantageous; future work should extend these findings using later releases of the ABCD Study^®^ data or other large brain imaging datasets that are in progress (e.g., Magnus et al. 2016).

In summary, this study documented age associations with adolescent structural brain morphometry at the ROI, vertex, and voxel level. Results replicated and extended previous findings related to adolescent brain development, and voxelwise Jacobian results illustrated distinct spatiotemporal patterns of volumetric changes in subcortical regions. This project also provides a shared resource (FEMA) that can be used by the public to run GAMMs within our fast and efficient computational framework. The tools developed for this project will serve as prototypes for future researchers interested in testing additional research questions at the vertex and voxel level without sacrificing model complexity.

## Acknowledgements

Data used in the preparation of this manuscript were obtained from the Adolescent Brain Cognitive Development^SM^ (ABCD) Study (https://abcdstudy.org), held in the NIH Brain Development Cohorts (NBDC) Data Hub. This is a multisite, longitudinal study designed to recruit more than 10,000 children age 9-10 and follow them over 10 years into early adulthood. The ABCD Study® is supported by the National Institutes of Health and additional federal partners under award numbers U01DA041048, U01DA050989, U01DA051016, U01DA041022, U01DA051018, U01DA051037, U01DA050987, U01DA041174, U01DA041106, U01DA041117, U01DA041028, U01DA041134, U01DA050988, U01DA051039, U01DA041156, U01DA041025, U01DA041120, U01DA051038, U01DA041148, U01DA041093, U01DA041089, U24DA041123, U24DA041147. A full list of supporters is available at https://abcdstudy.org/federal-partners.html. A listing of participating sites and a complete listing of the study investigators can be found at https://abcdstudy.org/consortium_members/. ABCD consortium investigators designed and implemented the study and/or provided data, but did not necessarily participate in the analysis or writing of this report. This manuscript reflects the views of the authors and may not reflect the opinions or views of the NIH or ABCD consortium investigators.

The ABCD data repository grows and changes over time. The ABCD data used in this report came from ABCD Study Data Release 6.0 (DOI: https://doi.org/10.82525/jy7n-g441). A complete list of DOIs can be found at https://nda.nih.gov/abcd/abcd-annual-releases.html.

The authors wish to thank the youth and families participating in the ABCD Study^®^ and all staff involved in data collection and curation. D. Smith was supported by Kavli Innovative Research Grant under award number 2022-2195 as well as a Ruth L. Kirschstein Individual Predoctoral NRSA award (5F30DA057078-02) for this project.

## Data availability

ABCD Study^®^ data release 6.0 is available for approved researchers in the NIH Brain Development Cohorts (NBDC) Data Hub (DOI: https://doi.org/10.82525/jy7n-g441).

## Code availability

The code to analyze the data and generate all figures of this manuscript is available on GitHub: https://github.com/dmysmith/age

