## Supplementary Information for "Age associations with cortical and subcortical brain structure in adolescents age 9-17"

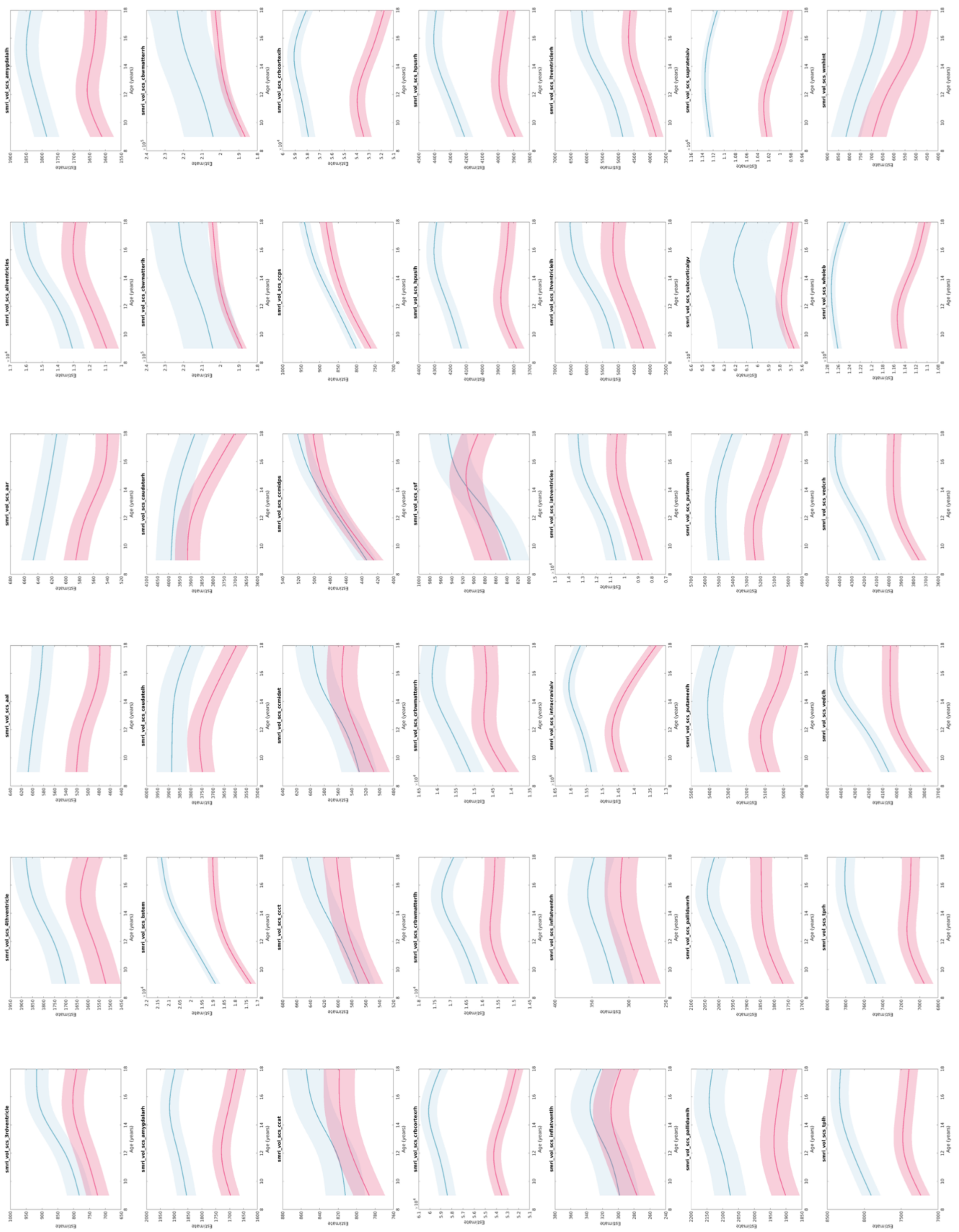

Supplementary Figure 1

Model-estimated subcortical volumes as a function of age. Estimates are shown in cubic millimeters on the y-axis. Error bars represent the analytic 95% confidence interval of the estimate. Variable names match those provided in the ABCD Data Dictionary for ease of replication. Pink line plot represents model estimated in females; blue represents estimates in males.

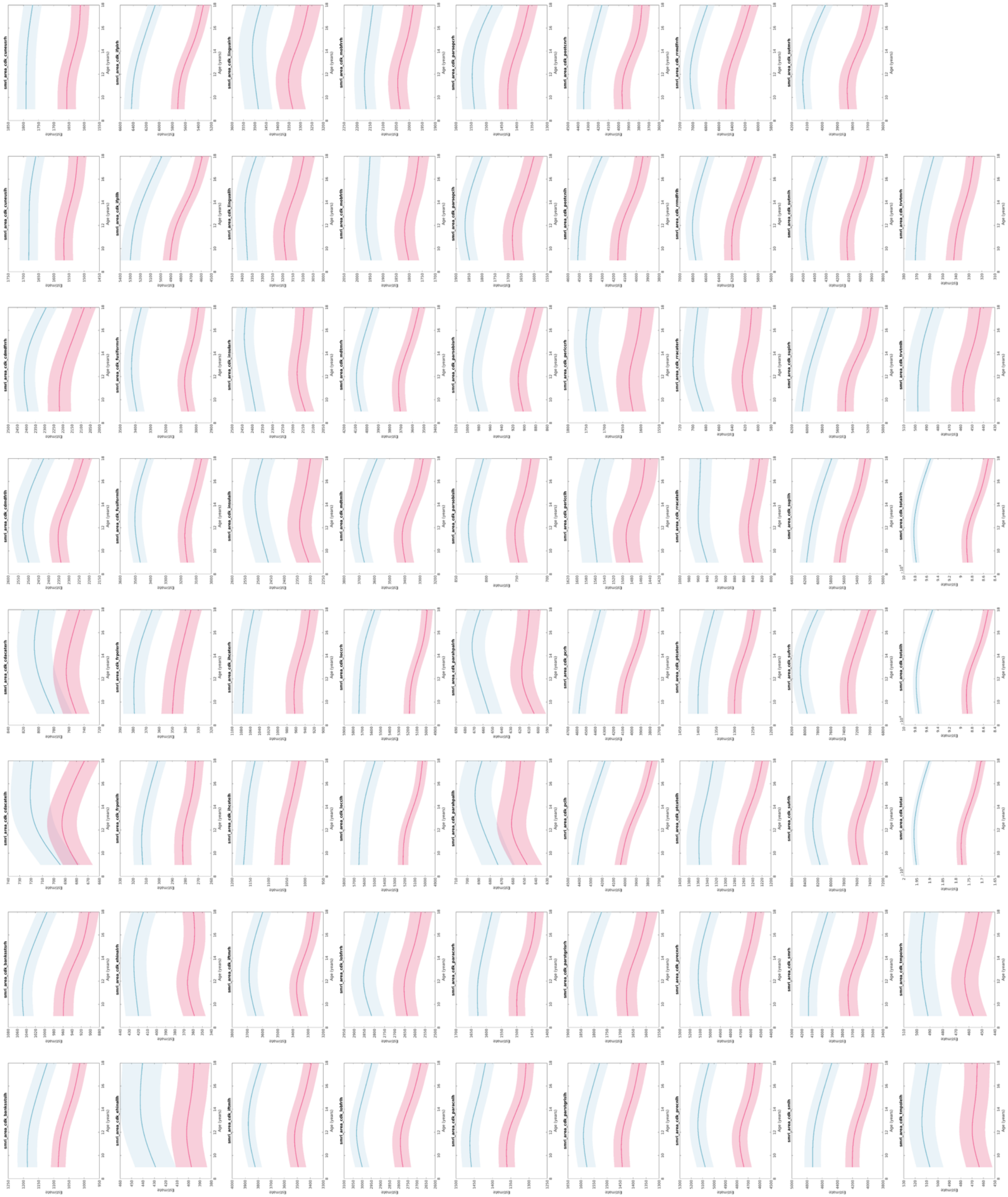

Supplementary Figure 2

Model-estimated cortical surface area (APARC parcellation) as a function of age. Estimates are shown in square millimeters on the y-axis. Error bars represent the analytic 95% confidence interval of the estimate. Variable names match those provided in the ABCD Data Dictionary for ease of replication. Pink line plot represents model estimated in females; blue represents estimates in males.

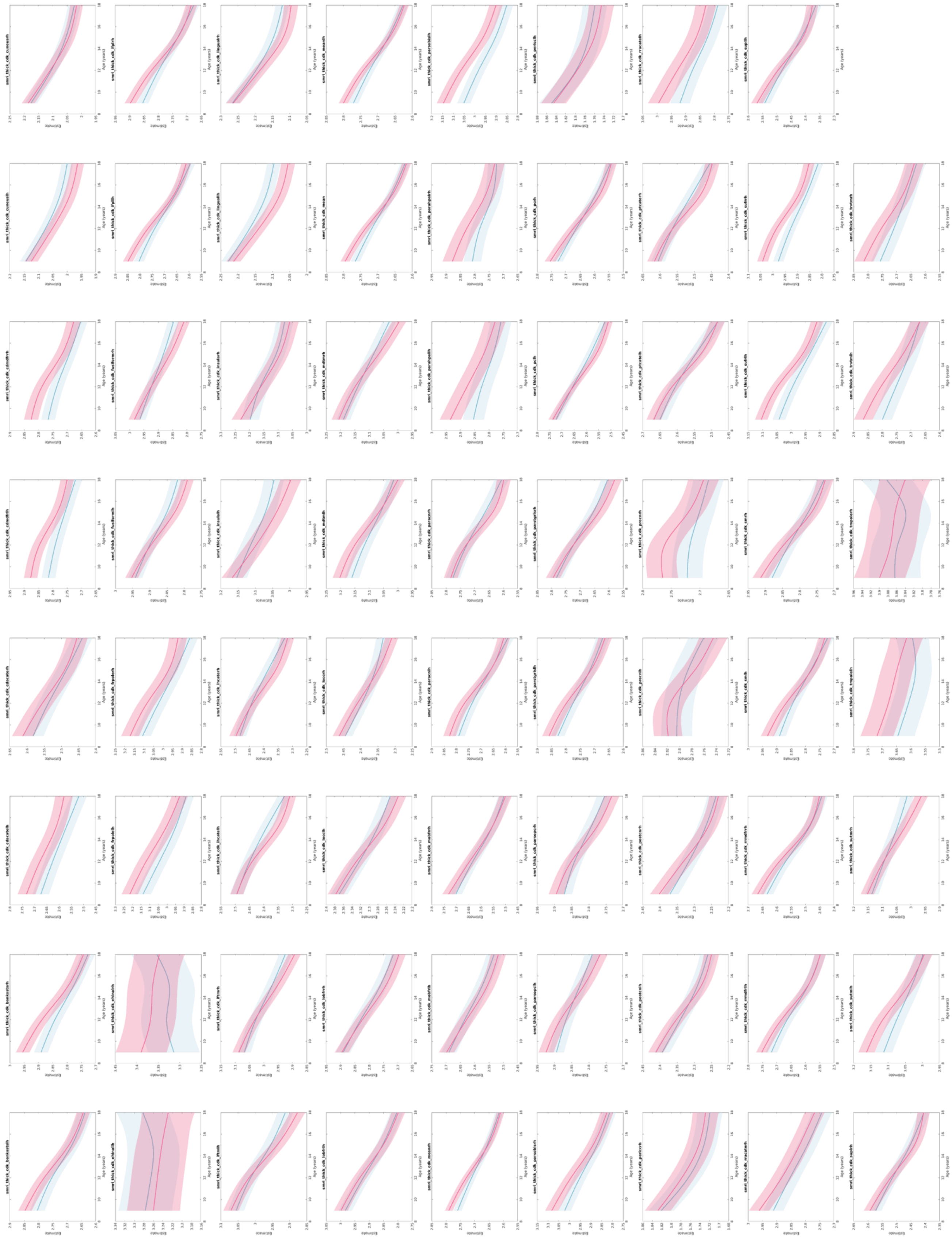

Supplementary Figure 3

Model-estimated cortical thickness (APARC parcellation) as a function of age. Estimates are shown in millimeters on the y-axis. Error bars represent the analytic 95% confidence interval of the estimate. Variable names match those provided in the ABCD Data Dictionary for ease of replication. Pink line plot represents model estimated in females; blue represents estimates in males.

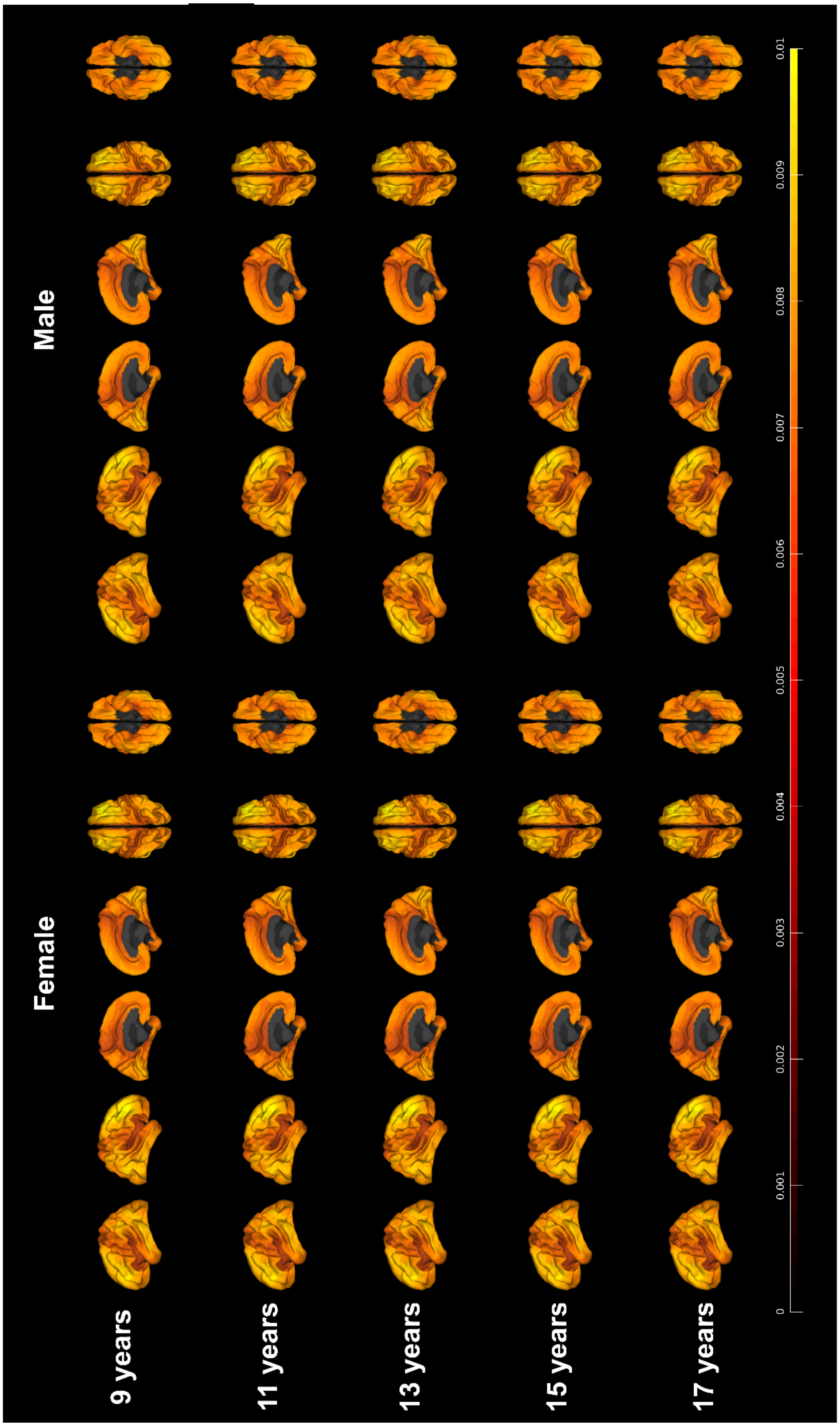

Supplementary Figure 4

Standard error of cortical surface area estimated at the vertexwise level, in units of square millimeters. Areas are estimated separately in females (left six columns) and males (right six columns).
